# Trajectory of rich club properties in structural brain networks

**DOI:** 10.1101/2021.11.16.468806

**Authors:** Levin Riedel, Martijn P. van den Heuvel, Sebastian Markett

## Abstract

Many organizational principles of structural brain networks are established before birth and undergo considerable developmental changes afterwards. These include the topologically central hub regions and a densely connected rich club. While several studies have mapped developmental trajectories of brain connectivity and brain network organization across childhood and adolescence, comparatively little is known about subsequent development over the course of the lifespan. Here, we present a cross-sectional analysis of structural brain network development in N = 8,066 participants aged 5 to 80 years. Across all brain regions, structural connectivity strength followed an ‘inverted-U’-shaped trajectory with vertex in the early 30s. Connectivity strength of hub regions showed a similar trajectory and the identity of hub regions remained stable across all age groups. While connectivity strength declined with advancing age, the organization of hub regions into a rich club did not only remain intact but became more pronounced, presumingly through a selected sparing of relevant connections from age-related connectivity loss. The stability of rich club organization in the face of overall age-related decline is consistent with a “first come, last served” model of neurodevelopment, where the first principles to develop are the last to decline with age. Rich club organization has been shown to be highly beneficial for communicability and higher cognition. A resilient rich club might thus be protective of a functional loss in late adulthood and represent a neural reserve to sustain cognitive functioning in the aging brain.

## 1 Introduction

The human brain is an intricate network whose complex wiring diagram can be reconstructed *in vivo* from magnetic resonance imaging (MRI) data and abstracted in a connectome network map (Park & Friston, 2013; Sporns, 2011; Hagmann, 2005; Hagmann et al., 2007; Sporns et al., 2005). The application of network analytics to such connectome maps has revealed that brain-wide topology of structural fiber connections follows several principles (Bullmore & Sporns, 2009): Across the brain, regions differ considerably in the number of their interconnections with other regions. Few regions claim the lion’s share of connections (Hagmann et al., 2008) and act as hubs in the brain network (van den Heuvel & Sporns, 2013). Hubs are multi- and transmodal regions that are topologically central in the network (Gong et al., 2009; Sporns et al., 2007), metabolically expensive (Collin et al., 2014), and involved in the integration of modular and segregated brain function (Bertolero et al., 2015; Cohen & D’Esposito, 2016; van den Heuvel & Sporns, 2013; Sporns, 2013). Highly connected brain regions tend to connect stronger to other highly connected regions than expected by their high number of connections alone, ultimately forming a rich club of densely interconnected brain regions that form the backbone for global brain communication (van den Heuvel et al., 2012; van den Heuvel & Sporns, 2011).

The human brain undergoes developmental changes across the lifespan (Sowell et al., 2004). While gray matter volume decreases non-linearly from childhood to old age, white matter volume and the integrity of fiber connections follow an ‘inverted U’ shaped trajectory with increases into mid-adulthood and a decline thereafter (Kochunov et al., 2012; Sowell et al., 2003). This raises the question whether topological features of brain network organization follow similar life span trajectories. Major organizational principles of the structural connectome such as network hubs and a rich club are already present as early as gestational week 30, suggesting that network formation occurs prenatally during the second trimester (Ball et al., 2014). During the third trimester, major maturation occurs on connections from the rich club to the rest of the connectome network (i.e. on so-called feeder connections), a pattern that continues across childhood (Wierenga et al., 2018). The overall tendency of high-degree nodes to connect preferably to other high degree nodes which gives rise to the rich club phenomenon, however, does not seem to change between childhood and adulthood (Grayson et al., 2014), even though the connectivity strength between rich club areas increases during adolescence (Baker et al., 2015). These findings are in line with a developmental model where qualitative principles are present from very early on and then develop quantitatively in two subsequent stages with connections of unimodal and peripheral brain regions maturing during childhood, and connections of multimodal and central hub regions maturing during adolescence (Wierenga et al., 2015). While network maturation during childhood and adolescence has received considerable attention (see also Hagmann et al., 2010), less is known about developmental trajectories across the adult life span. One report has found ‘inverted U’ shaped trajectories for nodal connectivity strength and efficiency (Zhao et al., 2015). Maturation of connections between hub regions peak earlier (i.e., late 20s/early 30s) than connectivity between hubs and the periphery (late 30s), resulting in a linear decrease of rich club organization across the life span. This finding stands in contrast to life span evidence on functional connectivity that suggests an ‘inverted U’ shaped trajectory (with peak at around age 40) for rich club organization (Cao et al., 2014). Given the paucity of life span data on connectome organization, the present report seeks to re-examine the life span trajectory of network hubs and the rich club in structural brain networks. We will go beyond previous studies by dramatically increasing the sample size and utilizing a large cross-sectional data set with structural connectomes of N = 8,066 participants aged 5 to 80 years. Specifically, we seek to map the life span trajectories of hub connectivity, the identity of hub regions according to data-driven criteria for hub definition, and rich club properties of the brain network.

## 2 Methods

### 2.1 Participants

We used openly available connectome data from the 10kin1day data set (van den Heuvel et al., 2019). This data set contains structural connectome data from N = 8,168 participants (n = 3824 females, n = 4339 males, n = 5 no gender specified) who are either classified as healthy controls or as patients with psychiatric or neurological illness (n = 4481 controls, n = 3668 patients, n = 19 no disease status specified). Patient status is given as binary category and no details on the precise diagnosis is given. The connectome dataset contains participants aged 0 to 90 years in 19 age groups (see table 1). Quality assurance and outlier removal have been performed by the curators of the data set. For our analysis, we additionally excluded the age groups 0, 0-5, 80-85, and 85-90 due to their small sample size, as well as the participants with unknown disease status or gender, resulting in a sample of N = 8,066 participants (n = 3776 females, n = 4290 males). The 10kin1day data set is the result of a three-day pop-up data processing event and contains jointly analyzed data from 42 different research groups. Informed written consent was obtained from all participants at each acquisition site and the protocols were approved by the local ethics committees at the independent research institutions.

**Table 1.** Age, gender and patient status distribution of the participants included in the 10kin1day dataset.

|  | Gender |  |  | Patient status |  |  |
| --- | --- | --- | --- | --- | --- | --- |
| Age group | Females | Males | Unknown | Controls | Patients | Unknown |
| 0 | 3 | 9 | 0 | 0 | 12 | 0 |
| 0 - 5 | 14 | 20 | 1 | 0 | 35 | 0 |
| 5 - 10 | 57 | 75 | 0 | 76 | 56 | 0 |
| 10 - 15 | 182 | 201 | 0 | 316 | 66 | 1 |
| 15 - 20 | 389 | 398 | 0 | 567 | 220 | 0 |
| 20 - 25 | 666 | 637 | 0 | 854 | 445 | 4 |
| 25 - 30 | 446 | 639 | 0 | 658 | 422 | 5 |
| 30 - 35 | 300 | 424 | 1 | 380 | 342 | 3 |
| 35 - 40 | 223 | 319 | 0 | 258 | 283 | 1 |
| 40 - 45 | 265 | 291 | 0 | 249 | 307 | 0 |
| 45 - 50 | 272 | 248 | 0 | 258 | 262 | 0 |
| 50 - 55 | 272 | 249 | 1 | 224 | 298 | 0 |
| 55 - 60 | 226 | 211 | 0 | 163 | 271 | 3 |
| 60 - 65 | 191 | 217 | 0 | 121 | 287 | 0 |
| 65 - 70 | 162 | 206 | 1 | 186 | 183 | 0 |
| 70 - 75 | 100 | 120 | 1 | 95 | 124 | 2 |
| 75 - 80 | 39 | 60 | 0 | 51 | 48 | 0 |
| 80 - 85 | 14 | 14 | 0 | 21 | 7 | 0 |
| 85 - 90 | 3 | 1 | 0 | 4 | 0 | 0 |

### 2.2 Data acquisition and processing

The 10kin1day data set includes imaging data from 42 different groups acquired on different scanners with varying field strength (1.5 and 3T) and acquisition parameters. Data were processed with a unified pipeline. Details on the processing pipeline and quality control are given in van den Heuvel et al. (2019). In brief, connectomes were assembled by first obtaining a cortical and subcortical gray matter parcellation from running T1-weighted structural images through Freesurfer (Fischl et al., 2004) and then collating the resulting parcellation with DTI data. Diffusion data were first corrected for susceptibility and eddy current distortions. Then each voxel’s main diffusion direction was obtained via robust tensor fitting. Large white matter pathways were formed by deterministic fiber tractography (Mori et al., 1999). Fiber streamlines were propagated along each voxel’s main diffusion direction after originating from eight seeds evenly distributed across each white matter voxel until a stopping criterion was met (hitting a voxel with FA <.1, a voxel outside the brain mask, or making a turn of > 45 degrees). A pair of regions from the gray matter parcellations was considered connected when both regions were touched by a reconstructed streamline. Connections were weighted with different metrics, of which we used three in the present report: the total number of streamlines (NOS) that touched both ROIs, mean fractional anisotropy (FA) of white matter voxels in reconstructed fiber tracts, and streamline-volume density (SVD, which is the number of streamlines normalized to the region volume). Gray matter regions were defined according to the Desikan-Killiany standard Freesurfer parcellation (aparc, Desikan et al., 2006) with 82 regions of interest. This resulted in three weighted (NOS, FA, SVD) and undirected connectome matrices for each individual. Unless stated otherwise, we present results from the NOS-weighted connectome matrices.

### 2.3 Network analysis

Network analyses were performed in MATLAB (MathWorks, version 20a), using the Brain Connectivity Toolbox (BCT, Rubinov & Sporns, 2010). The network analyses were used to first compare the average network connectivity between the different age groups. Then, the rich club property was analyzed for each individual and compared between age groups. Finally, based on the network connectivity and different criteria, including the rich club results, hub regions of each individual were defined, to also compare the average hub region connectivity between different age groups.

#### Connectivity analysis

All three different weights, number of streamlines (NOS), fractional anisotropy (FA), and streamline-volume density (SVD), were each averaged for all subjects individually, excluding non-existent connections. To account for differences in NOS and SVD weights based on brain region volume, regional volumes were averaged for all subjects. Also, the network density, maximum node degree and maximum connection weight (NOS) of each network were computed with their respective BCT functions (density_und.m, degrees_und.m). The calculations were done to later model connection weights and brain region volume for all age groups, while still accounting for inter-individual variability.

#### Rich club analysis

We followed standard procedures for rich club analysis (van den Heuvel et al., 2013). A network is said to have rich club properties when high degree nodes show a higher level of interconnectedness than expected from their high degree alone (van den Heuvel & Sporns, 2011), across a range of degree thresholds. The rich club regime was established as follows: We first computed the weighted rich club coefficient (using the BCT function rich_club_wu.m) across the full range of levels k from the network’s degree distribution (k=1, …, n). Because high degree nodes have a high likelihood to connect to other high degree nodes by chance alone, it is necessary to establish that the empirical level of interconnectedness exceeds the level of interconnectedness in random networks. We created 2,500 random networks per participant by reshuffling all connections in the matrix while preserving the degree distribution of the network (BCT function randmio_und.m). Each connection was rewired 10 times. At each level k, normalized rich club coefficients were then obtained by dividing empirical coefficients by the mean coefficient from all 2,500 iterations of the random network. We also used the distribution of random network coefficients to derive a p-value of the probability that the empirical rich club coefficient resulted from the non-selective high interconnectedness of high degree nodes. We corrected the false discovery rate (FDR) across the full range of p-values by applying the Benjamini & Hochberg (1995) procedure (Groppe, 2020). As a result, we obtained three curves across all levels k: a curve for the empirical rich club coefficient, a curve for the mean random rich club coefficient, and a curve for the normalized rich club coefficient. We derived our main outcome measures of individual rich club organization from these curves. We determined the rich club regime as the largest series of subsequent k, where the empirical rich club coefficient was larger than the rich club coefficient in 95% of all random networks (p-value < 0.05, FDR-corrected). We used the following algorithm to identify the rich club regime: We first selected the lowest and highest k-level with p <.05. If all interjacent p-values were also <.05, we defined the range between the two k-values as the rich club regime. If this was not the case, we applied the following: If only one single or two non-neighboring p-values within this range were >.05, we still considered the range as rich club regime (thus considering these datapoints as outliers). In case that two or more neighboring p-values exceeded the .05 threshold (i.e., cutting the range between the lowest and highest k-level with p<.05 in two or more), we assessed whether any of the ranges exceeded the other ones by a factor of 1.5. If this was the case, we assumed this range as the rich club regime. If no range was 1.5-times larger, we assumed the range with larger k values (i.e., at the upper end of the normalized rich club curve) as the rich club regime. If none of the criteria applied, we did not assume a valid continuous rich club regime for this participant. This was the case in 362 participants (~4.5%) who were excluded from further analysis. Once a valid rich club regime was established, we computed the following measures for each participant: We defined the length of the rich club regime as the difference between the upper and lower end of the rich club regime (if, for example, the empirical rich club coefficient exceeded the random rich club coefficient with p<.05, corrected, on all k-levels between k=12 and k=27, the length of the rich club regime was determined to be 15). A longer rich club regime might imply that more brain regions belong to the rich club. We also assessed this directly by determining how many nodes had a nodal degree equal to or larger than the first k-level belonging to the rich club regime. While these measures inform us on the size of the rich club in terms of its members, it does not give us information on the strength of the rich club effect, i.e., on how much the empirical rich club coefficient exceeds the rich club coefficient in comparable random networks. We therefore extracted the peak of the normalized rich club curve as a point estimate and also calculated the area under the normalized rich club curve above 1 with the trapezoidal method (trapz.m). The latter measure scales both with the number of rich club members and the strength of the rich club effect. Because the normalized rich club coefficient is a ratio between empirical and random coefficients, a separate analysis of the numerator and denominator can also be of interest. We therefore extracted additionally the areas under the empirical and random rich club curves (including the area below 1).

To distinguish the effect of changes in nodal degree or connection weights on the rich club results, the same analysis as described above was performed on binary connectome matrices. We used the respective BCT function (rich_club_bu.m) to compute binary rich club coefficients for each participant’s empirical network and for 2,500 permuted versions of the network. As the intention for this analysis was a direct control-comparison for the weighted rich club results, we based all outcome measures on the rich club regimes from the weighted analysis. We extracted the peak of the normalized binary rich club curve and the area under the normalized, empirical and random binary rich club curves for each individual.

#### Hub analysis

Hub definition is commonly based on different centrality-related network metrics and statistical criteria (van den Heuvel & Sporns, 2013). We defined hub participation according to five separate criteria: Brain regions were classified as hubs when they either belonged to the top 15% of the degree distribution (criterion a), to the top 15% of the strength (i.e. weighted degree) distribution (criterion b), when their nodal degree was equal to or larger than the nodal degree of the starting point of the rich club regime (criterion c), equal to or larger than the nodal degree of the peak of the normalized rich club curve (criterion d), or based on hub scores (criterion e). The hub score measure (criterion e) was composed of five centrality measures: nodal degree, betweenness centrality, nodal path length, between-module participation coefficient, and within-module degree z score. Nodal degree and betweenness centrality were calculated with their respective BCT functions (betweenness_wei.m, degrees_und.m). Between-module participation coefficient and within-module degree z score were calculated with BCT functions (participation_coef.m and module_degree_zscore.m) based on module parcellations for each combination of age group and disease status. Module parcellations were identified with the BCT function bct_community.m based on group connectomes that contained the average connection weights for all connections present in at least 60 % of the group’s participants. Nodal path length was calculated as the sum of the node’s distance to other nodes (as given by the BCT function distance_wei.m) divided by the number of all other nodes. Betweenness centrality and nodal path length used the connectivity-length matrix (as given by 1/connection weight) to represent higher connection weights as shorter paths. Brain regions present in the top 33 % of at least four out of five centrality measures were defined as hub regions according to hub scores ( van den Heuvel et al., 2015). We verified this hub-definition by a complementary approach that used the k-means algorithm (with k = 2) to partition all nodes into hub and non-hub regions based on all five centrality measures from the hub score measure (criterion e) (Markett et al., 2020).

We defined hubs in each individual brain network according to the criteria outlined above. For each hub definition, we then defined group level hubs as those twelve brain regions (i.e. ~15%) that were most consistently identified across participants. We derived hub definitions for each age group and for the whole sample. Pairwise similarity between different hub definitions was assessed across hub-criteria and across age groups with the Jakkard index (Steen et al., 2011). The similarity of the hub definitions for all age groups according to the hub score criterion as assessed by the Jakkard index was plotted as a heatmap in MATLAB. We used the hub definitions to compute the average connection weight of hub regions, the average gray matter volume of hub regions, and normalized versions of these measures by dividing by hub connection weights (and hub gray matter volume respectively) by the mean connection weight (gray matter volume) across the entire brain.

### 2.4 Statistical analysis

Statistical analysis was performed in R Studio (version 1.3.1056, R version 4.0.2). All lifespan changes were modelled with generalized additive mixed-effect models (GAMM; Lin & Zhang, 1999) using the gamm4 toolbox. GAMMs are based on linear mixed-effect models (gamm4 is based on the lme4 package) but with multiple sinusoidal base functions, whose number is automatically selected during the modelling process and is represented by the estimated degree of freedom (EDF) of each respective model. This allows the modeling of non-linear relationships without any *a priori* assumptions on the model type. Similar to linear mixed-effect models, GAMMs can include factorial variables and random effects. We modelled age nonparametrically, with gender and disease status as factorial variables. The age of each age group was set to the mean value of the range it covers, therefore representing the participants in each age groups at the same age. Research center and participants were included as a random term. To assess differences between gender and patient groups, we set up additional models differentiating all factors of the respective variable. The used model function was:

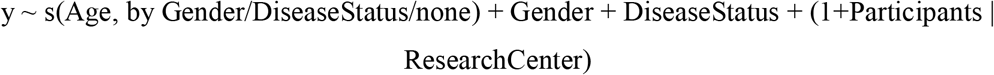

All models were plotted with the mgcv toolbox, including the 95 % confidence interval (CI) as shading. The fitted values and confidence intervals were extracted from the plotted models to assess peak values and their respective age value within the separate models.

### 2.5 Code and data availability

All data analyzed in the present report can be obtained upon request at dutchconnecomelab.nl. All analysis scripts will be made available on the open science framework upon publication.

## 3 Results

### 3.1 Rich club properties over the lifespan

A significant rich club regime was present in the connectomes of N = 7,704 participants (i.e. 95.5%). All subsequently reported analyses on rich club organization are focused on this group. We evaluated six summary measures from the rich club curve: The normalized rich club coefficient at the peak of the curve (i.e. the maximum difference between the empirical rich club coefficient and the null models), the length of the rich club regime (i.e. the range of the degree distribution for which the empirical rich club coefficient exceeded the rich club coefficient in the null models), the number of nodes with a nodal degree equal to or larger than the nodal degree of the starting point of the rich club regime (i.e. the number of nodes which qualify for the rich club), the area under the normalized rich club curve above 1 (i.e. the combination of the rich club regime length and peak value), and the area under the empirical and random rich club curves (i.e. the properties of the empirical and random networks defining the rich club property). Fitted values for all six measures across the lifespan and the control analyses with binary connectome matrices are shown in figure 1.

**Figure 1:**
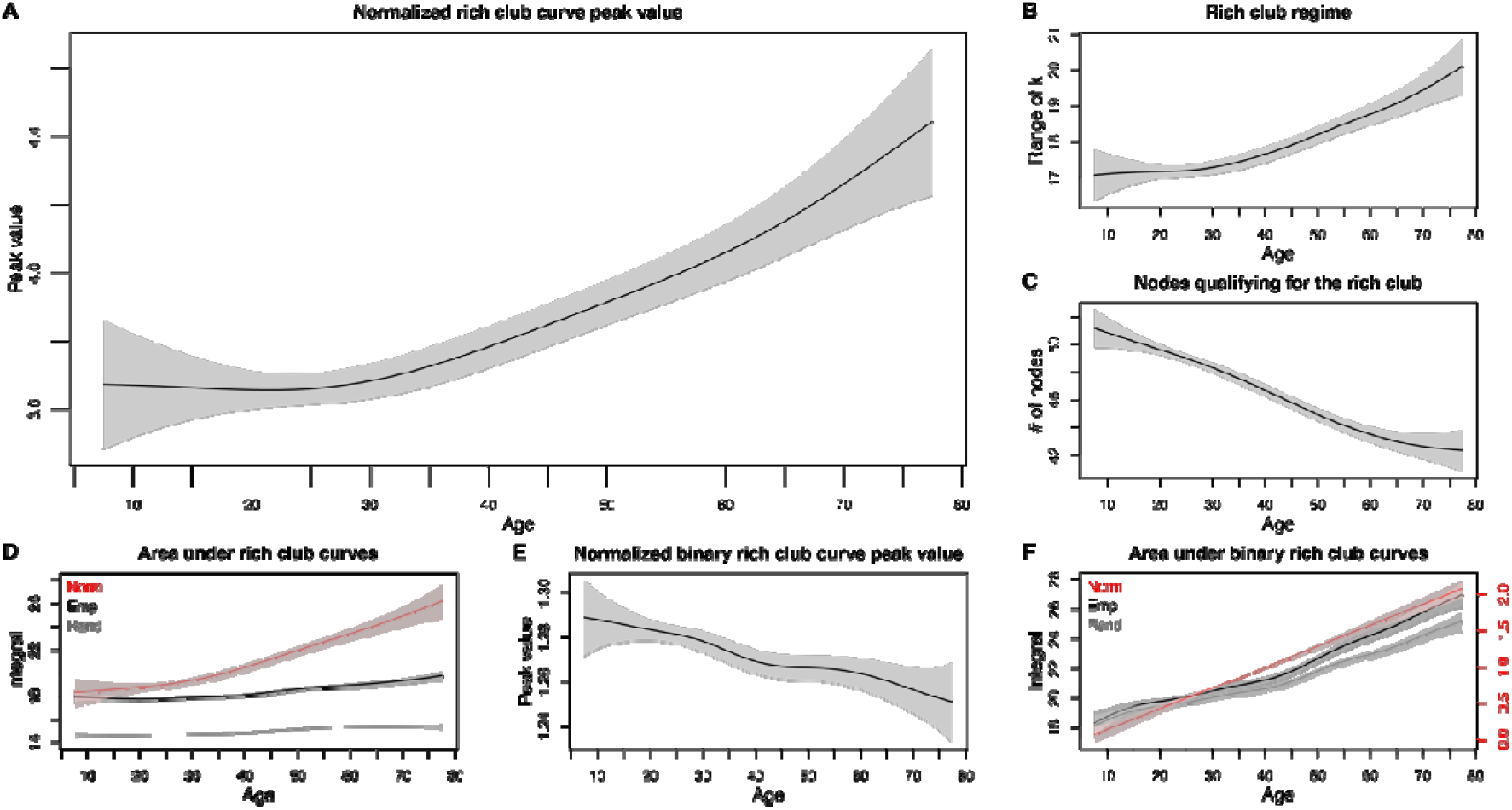
Rich club organization across the lifespan. Rich club organization as indexed by the maximal normalized weighted rich club coefficient increases with age (A). The rich club regime becomes longer with age, indicating a higher variance in nodal degree across rich club members (B) while the size of the rich club in terms of implicated brain regions decreases with age (C). The overall increase in rich club organization (area under the normalized rich club curve) is likely to result from a selective sparing of connections between rich club members as indicated by a less steep increase of rich club organization in the random null models (D). The analysis of binary versions of the networks (E and F) indicates that the observed changes in rich club organization are not reflected in the presence or absence of connections but rather reflect quantitative changes in connectivity strength. The color coding distinguishes the area under the normalized rich club curve (red) and areas under the rich club curve for the empirical networks (black) and the random null models (gray). Plotted are fitted values from GAMMs with 95% confidence intervals as shading.

GAMM-modeling revealed an increase in rich club organization across the lifespan: The peak of the normalized rich club curve increased with age (Figure 1A; EDF = 3.21, F = 18.92, p = 1.39e-12), i.e. the difference between empirical rich club organization and rich club organization in random null networks became more pronounced. At its peak, the empirical rich club coefficient exceeded the null models’ mean coefficient by a factor of 3.66 ± .05 in childhood and by 4.44 ± .22 in late adulthood. With increasing age, the rich club regime also became more widespread (Figure 1B; EDF = 3.684, F = 19.62, p = 1.76e-14), i.e., regions within the rich club would differ more in their nodal degree. The length of the rich club regime increased from 17.1 ± .72 (mean ± 95 % CI) in childhood to 20.1 ± .79 in late adulthood. Contrary to the increasing rich club regime, the number of nodes qualifying for the rich club decreased with age (Figure 1C; EDF = 3.68, F = 61.62, p < 2e-16), i.e., fewer nodes within the brain network show higher interconnectedness to other high degree nodes than expected by their nodal degree alone with increasing age. The number of nodes with a nodal degree equal to or larger than the nodal degree at the starting point of the rich club regime decreased from 51.2 ± 1.4 in childhood to 42.3 ± 1.52 in late adulthood. Of note, the observed increased difference for the rich club regime and peak value was also present in the whole area under the curve (Figure 1D, red; EDF = 2.96, F = 45.95, p < 2e-16), with an increase in the area under the curve above 1 from 18.3 ± 1.27 in childhood to 26.1 ± 1.46 in late adulthood, indicating that the rich club becomes relatively richer. The area under the empirical rich club curve (Figure 1D, black; EDF = 7.876, F = 25.32, p < 2e-16) and the random rich club curve (Figure 1D, gray; EDF = 4.739, F = 26.37, p < 2e-16) also increased with increasing age, but at different rates. The area under the empirical rich club curve increased from 17.7 ± .11 at age 20 to 19.7 ± .43 in late adulthood, while the area under the average random rich club curve only increased from 14.6 ± .09 at age 17 to 15.4 ± .13 in late adulthood, resulting in an increase of the overall rich club property with increasing age.

As network metrics are known to relate to network density, we modelled average network density and maximum nodal degree over the lifespan. Here, no age effect was detectable for the average network density (Supplementary figure 1A; EDF = 1.784, F = 2.874, p = .0608) but the maximum nodal degree increased across the lifespan (Supplementary figure 1B; EDF = 6.45, F = 67.23, p < 2e-16). To control for the increase in maximum nodal degree across the lifespan we performed the rich club analyses also on binary connectome matrices. The peak of the normalized binary rich club curve actually decreased with age (Figure 1E; EDF = 5.093, F = 5.583, p = 3.39e-5), but the area under the normalized (Figure 1F, red; EDF = 1.009, F = 316.4, p < 2e-16), empirical (Figure 1F, black; EDF = 6.456, F = 97.93, p < 2e-16) and random (Figure 1F, gray; EDF = 7.004, F = 82.65, p < 2e-16) binary rich club curves increased. It must be noted though, that the overall quantity of the normalized rich club curve peak value (childhood: 1.29 ± .02; late adulthood: 1.25 ± .02) and the area under the normalized binary rich club curve (childhood: .06 ± .09; late adulthood: 2.08 ± .13) was very low. The latter is also represented by the very similar trajectory and quantity of the areas under the empirical and random binary rich club curves and indicates little to no rich club property observed using the binary connectome matrices. Taken together, these results show that the observed increase in the peak value of the normalized weighted rich club curve is not driven by the increase in the maximum nodal degree over the lifespan, as this increase would have been observed using binary connectome matrices, whereas the increase in the area under the different rich club curves and the rich club regime might in part by driven by this increase. From these results we conclude that rich club organization in structural brain networks is preserved over the lifespan, which is in contrast to previous reports suggesting a decrease or an ‘inverted U’ shaped pattern.

### 3.2 Hub regions over the lifespan

We computed hub scores to classify brain regions into likely hubs and non-hubs for each individual in the data set (see methods). At the group level, we defined hubs as those twelve brain regions (~15%) that were most consistently classified as hubs at the individual level. This resulted in the following hub regions: left and right thalamus, left and right putamen, left and right superior frontal as well as superior parietal gyrus, left and right precuneus, and left and right insula (see table 2). Alternative methods for individual hub detection via nodal degree, the starting point of the rich club regime, or the peak value of the normalized rich club curve resulted in the same hub vs. non-hub partition (all pairwise Jakkard indices J =1). Only the hub definitions via the strength (weighted degree) distribution and via k-means clustering of the hub score centrality measures resulted in a slightly different group-level hub assignment comparing to all other criteria (J =.5, equivalent to 4/12 different hubs), but were more similar comparing to each other (J = .71, equivalent to 2/12 different hubs). Given the largely consistent results across partitioning approaches, we decided to retain the hub definition based on hub scores for further analyses. The partition of brain regions into hubs and non-hubs was highly similar across age groups (see figure 2), resulting in an identical hub vs. non-hub partition for almost all age groups (with hub regions as listed above; all pairwise Jakkard indices J = 1). The only differences were observed in the age groups 5-10, 10-15, 15-20, and 75-80 (J = 0.85, equivalent to 1/12 different hubs, or J = .71, equivalent to 2/12 different hubs).

**Table 2.** Percent nodal participation in the top 15 % hub region for all participants according to the hub scores measure. Bold marks the top 12 nodes (~15 %), which are considered the group-level hubs.

| Number | Node | Hub participation [%] |
| --- | --- | --- |
| **1** | **Left putamen** | **99,58** |
| **2** | **Right putamen** | **98,95** |
| **3** | **Left superiorfrontal** | **95,96** |
| **4** | **Left thalamus** | **93,44** |
| **5** | **Right superiorfrontal** | **92,86** |
| **6** | **Left superiorparietal** | **90,66** |
| **7** | **Right superiorparietal** | **82,64** |
| **8** | **Right thalamus** | **77,52** |
| **9** | **Right precuneus** | **76,82** |
| **10** | **Left precuneus** | **75,49** |
| **11** | **Left insula** | **66,49** |
| **12** | **Right insula** | **65,87** |
| 13 | Left caudate | 56,30 |
| 14 | Left rostralmiddlefrontal | 52,83 |
| 15 | Left precentral | 47,37 |
| 16 | Right rostralmiddlefrontal | 42,98 |
| 17 | Right caudate | 33,35 |
| 18 | Left pallidum | 31,47 |
| 19 | Right precentral | 25,84 |
| 20 | Left hippocampus | 21,83 |
| 21 | Right pallidum | 17,99 |
| 22 | Left superiortemporal | 15,77 |
| 23 | Left inferiorparietal | 13,95 |
| 24 | Right hippocampus | 13,33 |
| 25 | Left lateralorbitofrontal | 11,03 |
| 26 | Left isthmuscingulate | 10,96 |
| 27 | Right superiortemporal | 8,78 |
| 28 | Right isthmuscingulate | 8,77 |
| 29 | Left postcentral | 7,96 |
| 30 | Left lateraloccipital | 7,85 |
| 31 | Right inferiorparietal | 5,83 |
| 32 | Right lateralorbitofrontal | 4,81 |
| 33 | Left medialorbitofrontal | 4,04 |
| 34 | Left middletemporal | 3,00 |
| 35 | Right posteriorcingulate | 2,83 |
| 36 | Left posteriorcingulate | 2,58 |
| 37 | Right lateraloccipital | 2,16 |
| 38 | Right postcentral | 1,38 |
| 39 | Right caudalanteriorcingulate | 1,15 |
| 40 | Right rostralanteriorcingulate | 1,14 |
| 41 | Right medialorbitofrontal | 1,12 |
| 42 | Right middletemporal | 1,12 |
| 43 | Left rostralanteriorcingulate | 1,03 |
| 44 | Left inferior temporal | 0,86 |
| 45 | Left fusiform | 0,82 |
| 46 | Right amygdala | 0,64 |
| 47 | Right fusiform | 0,58 |
| 48 | Left caudalanteriorcingulate | 0,56 |
| 49 | Right frontalpole | 0,56 |
| 50 | Right inferior temporal | 0,52 |
| 51 | Right lingual | 0,36 |
| 52 | Left lingual | 0,33 |
| 53 | Left frontalpole | 0,30 |
| 54 | Left amygdala | 0,24 |
| 55 | Right paracentral | 0,22 |
| 56 | Right pericalcarine | 0,20 |
| 57 | Left cuneus | 0,19 |
| 58 | Left pericalcarine | 0,19 |
| 59 | Left temporalpole | 0,19 |
| 60 | Right cuneus | 0,19 |
| 61 | Left paracentral | 0,17 |
| 62 | Right temporalpole | 0,17 |
| 63 | Left parstriangularis | 0,16 |
| 64 | Left caudalmiddlefrontal | 0,10 |
| 65 | Right parsorbitalis | 0,07 |
| 66 | Right parstriangularis | 0,07 |
| 67 | Right accumbens area | 0,06 |
| 68 | Left supramarginal | 0,04 |
| 69 | Right supramarginal | 0,04 |
| 70 | Left accumbens area | 0,02 |
| 71 | Left bankssts | 0,01 |
| 72 | Left parsopercularis | 0,01 |
| 73 | Right caudalmiddlefrontal | 0,01 |
| 74 | Left entorhinal | 0 |
| 75 | Left parahippocampal | 0 |
| 76 | Left parsorbitalis | 0 |
| 77 | Left transversetemporal | 0 |
| 78 | Right bankssts | 0 |
| 79 | Right entorhinal | 0 |
| 80 | Right parahippocampal | 0 |
| 81 | Right parsopercularis | 0 |
| 82 | Right transversetemporal | 0 |

**Figure 2:**
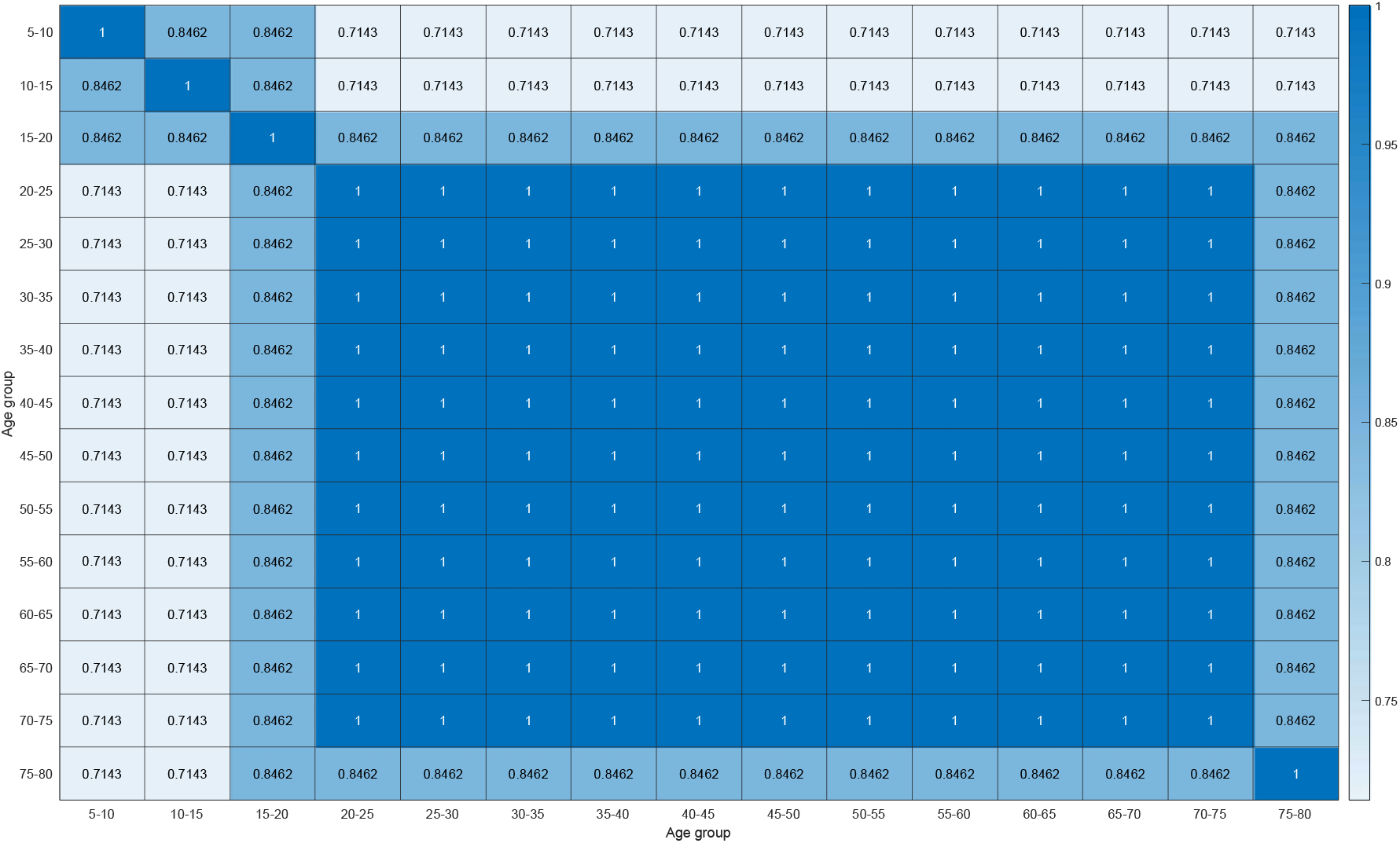
Similarity (expressed by the pairwise Jakkard index) of hub assignments (based on hub scores) between age groups.

### 3.3 Nodal and hub connectivity across the lifespan

We computed average connectivity in all individuals by averaging connection weights across all nodes or across all twelve hubs (see methods). Non-linear relationships between connectivity and participant’s age were modelled with GAMMs for 8,066 participants between the ages of five to 80 years.

Average connectivity across brain regions varied significantly with age (Figure 3A; EDF = 7.424, F = 45, p < 2e-16) and followed an ‘inverted U’ shaped trajectory across the lifespan. Average connectivity increased slightly from age 5 onwards, peaked at age 32, and showed a steep decline afterwards. We observed a similar trajectory for hub connectivity (Figure 3B; EDF = 4.961, F = 72.16, p < 2e-16) with a peak value at age 27. The direct comparison of hub connectivity and average connectivity indicated life span changes (Figure 3C; EDF = 6.537, F = 40.46, p < 2e-16): The relationship between hub- and average connectivity slightly increases between ages 5 and 22, with 2.75 ± .02-fold higher connectivity for hub regions at age 22. From 22 years onwards, the ratio between hub- and average connectivity decreased until age 67 to 2.37 ± 0.05-fold higher hub connectivity and again increased slightly afterwards. Please note that the decreasing ratio reflects relative changes in the connectivity of hubs vs. average connectivity and could be a consequence of the earlier apex of hub connectivity observable in the present data and as reported in previous work (Zhao et al., 2015). Unsurprisingly, the maximum connection weight demonstrated a very similar trajectory across the lifespan (Supplementary figure 1C; EDF = 6.402, F = 12.09, p = 4.77e-14).

**Figure 3:**
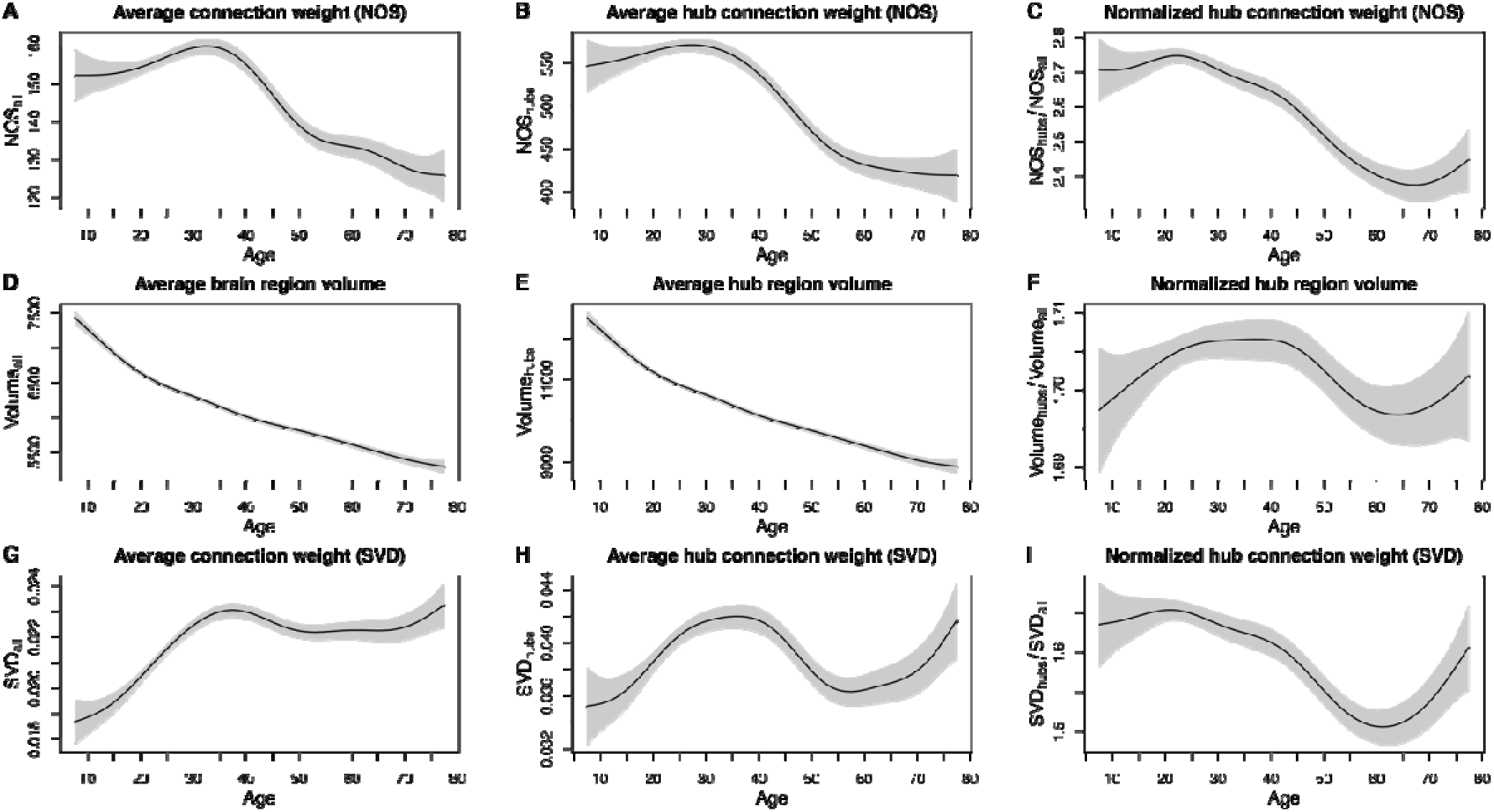
Average nodal and hub properties across the life span. Panels in rows correspond to: NOS-connection weights (row 1, A-C), regional gray matter volume (row 2, D-F), and SVD-weighted connection weights (row 3, G-I). Panels in columns refer to averages across all brain regions (column 1), averages across all hub regions (column 2), and averages of hub regions relative to all brain regions (column 3). Plotted are fitted values from GAMMs with 95% confidence intervals shaded in gray.

As larger brain regions are more likely to be touched by more reconstructed streamlines, it is necessary to evaluate connectivity changes from the perspective of changes in regional gray matter volumes. We found regional brain volume to decrease across the life span (Figures 3D-E; all regions: EDF = 7.503, F = 369.9, p < 2e-16; hub regions: EDF = 7.645, F = 359.4, p < 2e-16). The direct contrast of hub vs. whole brain regional volumes revealed small life span changes of the ratio (Figure 3F; EDF = 6.12, F = 3.769, p = 8.81e-4) that followed an ‘inverted U’ shaped trajectory. The fitted values, however, showed almost no difference with a minimum value of 1.7 ± .004 and a maximum value of 1.71 ± .003, indicating little evidence for relative changes in hub regional volumes.

Given the life span changes of regional brain volume and the confound of the NOS measure with regional brain volume, we further modelled life span trajectories of SVD-weighted connectivity. SVD-weighted connectivity followed an ‘inverted U’ shaped trajectory similar to NOS-weighted connectivity. Both average and hub connectivity, however, had a more pronounced increase with a peak at age 37 (average connectivity) and age 36 (hub connectivity). After this, average connectivity decreased moderately and remained at a relatively high level, while hub connectivity showed a more rapid decline. This was also reflected in a decrease in the ratio between hub and average connectivity from 1.65 ± .01 at age 21 to 1.51 ± .02 at age 61. All three measures (average connectivity, hub connectivity, ratio) increased again in late adulthood, with the average connectivity increasing further than its original peak at age 37 (Figures 3G-I; all regions: EDF=6.477, F = 31.88, p < 2e-16; hub regions: EDF = 7.271, F = 13.29, p < 2e-16; ratio: EDF=6.499, F = 20.38, p < 2e-16).

All hub analyses reported in this paragraph were based on the twelve brain regions that were most consistently identified as hubs across the entire sample. Because of subtle differences in hub regions in the age groups below 20 and above 75 (see figure 2), we explored two alternative hub partitions: Treating only those nine brain regions as hubs that were identified as hubs in each age group and using a group-specific hub definition of those twelve regions most consistently identified as hubs in the respective age groups lead to highly similar results. Please see the supplementary figure 2 for details.

### 3.4 Further analyses

All analyses reported above were statistically controlled for participants’ gender and patient status and for different study sites. We document fitted GAMM models for interactions between age and gender, and between age and patient status in the supplementary material (see supplementary figures 3-7). If not stated otherwise, all analyses reported above were based on weighted structural networks with NOS as connection weights. We document GAMM models for an alternative connection weight (i.e., FA) in the supplementary material (see supplementary figure 8).

## 4 Discussion

We present a cross-sectional analysis of life span trajectories of structural brain networks in N = 8,066 individuals aged 5 to 80. Our main findings are: 1) Structural connectivity across brain areas in general, and of highly connected hub regions in particular, follows an ‘inverted U’ shaped trajectory with an increase until middle adulthood and a decline afterwards, 2) regional gray matter volume decreases with age, and 3) rich club organization of the structural connectome is conserved across the life span. While the first two observations are confirmatory findings for previous literatures, the finding of conserved rich club organization is a novel discovery with implications for healthy brain aging, and neurological as well as cognitive reserve.

### Life-span trajectories in structural brain networks

The ‘inverted U’ shaped trajectory in structural connectivity confirms previous reports that either showed a similar trajectory across the life span (Kochunov et al., 2012; Zhao et al., 2015), or revealed consistent changes across selected age ranges such as increase in structural connectivity across childhood and adolescence (Hagman et al., 2008; Baker et al., 2015; Wierenga et al., 2015), or decrease in measures of white matter integrity from middle to late adulthood (Burzynska et al., 2010; Gong et al., 2009; Otte et al., 2015). The observed decrease in gray matter volumes is consistent with a large body of literature that reports such structural decline across the life-span (see Sowell et al., 2003, and Sowell et al., 2004, for review). Structural rich club organization has been shown to increase during childhood and adolescence (Baker et al., 2015; Wierunga et al., 2018), which is consistent with the present findings. One previous life span study, however, has described decreasing rich club organization in structural brain networks across the life span, an opposite pattern to the present finding, while reporting a similar ‘inverted U’ shaped trajectory for structural connectivity of hub regions (Zhao et al., 2015). This study, however, did only evaluate the normalized rich club coefficient at one statistically defined degree-level. The present findings provide a more detailed analysis by integrating rich club coefficients across the entire rich club regime derived from the normalized rich club curve.

Many network neuroscience studies define the rich club as a group of highly interconnected hub regions. Rich clubs, however, are not necessarily a nodal property of a few highly connected brain regions. Rather, the rich club reflects the organizational principle of the entire network that nodes prefer connecting to other nodes of equal or higher degree (van den Heuvel & Sporns, 2011). We found that rich club organization of the network as a whole is not only preserved but becomes even more pronounced in late adulthood. At first glance, this might appear at odds with the observed connectivity decrease of the most pronounced hub regions which started even earlier than the decrease in average connectivity across all nodes, as visible in the declining ratio of hub-over average connectivity (see figure 3C). The normalized rich club coefficient which quantifies rich club organization, however, relies on a within-subject comparison of empirical rich club connectivity with random network null models. The null networks are obtained by randomly shuffling edges while preserving the strength distribution of network nodes and are therefore similarly affected by changes in average connectivity. Our main finding of stronger rich club organization in older age is therefore a likely result from a targeted sparing of relevant connections of rich club members. Support for this explanation comes from the separate modeling of the empirical and random rich club curve across age: The increase of the empirical rich club curve is steeper than the random curve which leads to higher normalized rich club coefficients (see figure 1D). A second explanation is the decreasing number of nodes qualifying for club membership (figure 1C): Rich club coefficients become larger when fewer nodes are retained at a given threshold k, a pattern that also contributes to preserved rich club organization in Alzheimer’s disease (Daianu et al., 2013). It is important to note, however, that we only found an age-related increase in rich club organization when analyzing weighted networks. No such effect was observable in binary versions of the networks that only distinguished whether regions were connected by reconstructed fiber tracts but did not contain information on connectivity strength. The age-related increase in rich club organization is thus mainly reflected in a change of connectivity strength rather than in a qualitative remodeling of the brain network which would imply a loss of existing or a (biologically implausible) creation of new connections.

### Development of rich club organization - First come, served last?

The process of aging has been described as reversed ontology, where the last systems to mature are the first to decline. This observation of retrogenesis has been discussed in the context of dementia (Reisberg et al., 1999) but also regarding normal aging (Tamnes et al., 2013; Toga et al., 2006), together with the implicit assumption that the higher plasticity of late-maturing structures leaves them more vulnerable to degeneration. When also considering previous findings on rich club organization across childhood, the current observation that rich club organization is not only retained but enhanced in aging aligns with the concept of retrogenesis and suggests that rich club organization develops in a “first come, served last” principle across the lifespan. Rich club organization in structural brain networks has been observed as early as gestational week 30, suggesting that relevant connections are among the first that are created in the developing brain (Ball et al., 2014). Rich club organization remains stable between child- and adulthood (Grayson et al., 2014) and, according to our present finding, increases with advancing age, presumingly due to a targeted sparing of relevant connections from age-related decline. It remains open, however, if the persistence of rich club organization from the prenatal period to elderliness is supported by the same fiber connections and is thus a systems-level consequence of local developmental trajectories, or whether the organizational principle of the brain network is preserved through a systems-level reorganization.

While the current investigation looked solely into anatomical principles of structural brain network development, the rich club finding may still have important implications for brain function and cognitive aging. Higher rich club organization in the aging connectome could reflect a form of neural reserve or compensation to maintain function (Fornito et al., 2015). It has, for instance, been shown that stronger rich club organization in middle and late adulthood relates to better performance in cognitive domains (Baggio et al., 2015). The rich club provides a communication backbone which is relevant for the integration of segregated functional networks (de Reus & van den Heuvel, 2014; van den Heuvel & Sporns, 2013; van den Heuvel et al., 2012). The functional brain network seems to become less modular and more segregated in aging (Cao et al., 2014; Geerligs et al., 2015), and it has been suggested that the modular reorganization of the brain network could reflect compensatory efforts to maintain function in old age (Song et al., 2014). This could be attributable to the stronger rich club organization of the aging brain. The distinction between nodal changes and network-level changes has also been noted regarding network efficiency: Local efficiency, i.e. the inverse of the average shortest path of one node to its neighbors, declines with age, while global efficiency, i.e. the inverse of the average shortest path in the entire network, is typically unaffected (Cao et al., 2014; Geerligs et al., 2015; Song et al., 2014). From a methodological perspective, it is important to note that we did not observe age-related changes in network density. Network density can have marked influences on network metrics in brain networks (van Wijk et al., 2010) and needs to be accounted for when comparing different groups (van den Heuvel et al., 2017). Unfortunately, our present data set did not include cognitive or other function-related outcome measures. We can therefore only speculate whether preserved or increased rich club organization in the aging brain comes with functional benefits such as slower cognitive aging or a higher overall-functionality. This, however, would be an exciting prospect.

### Methodological considerations

Both linear and non-linear life span trajectories for brain development have been reported in the literature (Faghiri et al., 2019; Ziegler et al., 2012). Trajectories have often been described to follow an ‘inverted U’, but this does not necessarily imply strict quadratic development and it is generally recommended against parametric statistical models when modeling brain development as a function of age (Fjell et al., 2010). We addressed this issue by adapting generalized additive mixed-effect modeling (Wood & Scheipl, 2017). GAMMs are well suited to model life span brain trajectories (Sørensen et al., 2021; Walhovd et al., 2016) because they do not enforce parametric representations of age-brain relationships while additionally allowing fixed and random linear effect structures to control for possible confounds such as sex, patient status, and study site.

Through the 10kin1day data set, we were able to utilize the largest openly available structural connectome data set and to increase the sample size substantially in comparison to previous studies. The resulting increase in statistical power addresses a crucial methodological issue in neuroimaging (Button et al., 2013), particularly in the field of individual differences (Dubois & Adolphs, 2016). Achieving such sample size is virtually impossible without curated data from large consortia (Miller et al., 2016; Thompson et al., 2020; Van Essen et al., 2013), or open data sharing (Poldrack & Gorgolewski, 2014; Poline et al., 2012), an approach we benefited from in the present study. Through collaborative data sharing and unified preprocessing, the 10kin1day data set adapts an approach that has previously been successfully applied to functional brain networks (Bellec et al., 2017; Biswal et al., 2010). Connectome matrices from 10kin1day data show high correspondence with connectome matrices from the Human Connectome Project (HCP) which provides strong support for the validity of data (van den Heuvel et al., 2019). Nevertheless, combining imaging data from several centers comes of course with several challenges that cannot be fully compensated by the increase in sample size: Participants were scanned at different centers with different acquisition protocols and at different field strength. We were only able to control statistically for different study sites and cannot exclude the possibility of selection biases regarding different age groups at different centers. The apparent increase in connectivity in SVD-weighted connectomes after age 60, for instance, is likely a selection bias towards higher functional status in older participants, which is a common problem with convenience samples in aging research (Hultsch et al., 2002). Furthermore, the data set includes patient and non-patient data and no further information on specific diagnoses or assessments is given. We decided to include all participants and to control statistically for patient status. We are optimistic that the observed trajectories apply to both healthy people and people with neurological or psychiatric disorders. A more careful perspective on different conditions, however, would have been desirable. Lastly, the age variable was only available in bins (i.e., in age groups spanning five years). While this is an important measure towards protecting participants’ identity in a shared and widely accessible data set, it is of course a short coming when addressing developmental research questions. Lastly, we need to emphasize that our results rely on a cross-sectional comparison which cannot exclude cohort effects and does not allow for inferences on causality or the succession of age-associated alterations in network organization. We would therefore consider our current findings as tentative and encourage replication, for instance with data from the HCP lifespan project (Bookheimer et al., 2019) or in combined cross-sectional and longitudinal designs (Fotenos et al., 2005).

### Brain aging

The present study adds to the literature on changes in brain structure and organization across the life span. While early and late-life development of the structural connectome have been studied in isolation before, only few studies have taken a life-span perspective from early childhood to late adulthood in one data set. Taking a life-span perspective on brain development, however, is crucial as the pattern of early-life development and late-life decline shows a certain degree of overlap (Tamnes et al., 2013) and early-life development seems to set the stage for late-life decline (Deary et al., 2006; Walhovd et al., 2016). While previous studies also suggest an ‘inverted U’ shaped trajectory of white matter connectivity (Kochunov et al., 2012; Zhao et al., 2015), the apex of the developmental curves was found to vary between the late 20s and early 30s. Our data from a larger sample suggest that decline in average connectivity and hub connectivity might even begin a few years later. Future work will want to address the question how such decline is triggered, if there are ways to slow it down, and at which point possible interventions would be most effective. The biological aging process is characterized by a build-up of damage and limits of somatic maintenance throughout adulthood (Ferrucci et al., 2020; Hamczyk et al., 2020; Kirkwood, 2005). This leaves aging as a major risk factor for several prevalent conditions (Niccoli & Partridge, 2012), including neurodegenerative disorders whose incidence increase dramatically in older age (Querfurth & LaFerla, 2010). Timed interventions at an age where no functional loss has occurred as of yet might be most effective towards counteracting developmental decline and prolonging health over the lifespan (Ferrucci et al., 2020). It is an important observation in life-span research that age-related loss of function is highly individual (Lindenberger, 2014). While some individuals seem to age early, others maintain a high level of functioning into very old age. Several studies have therefore followed the approach to estimate individual brain age, which is thought to reflect the aging process better than chronological age (Cole & Franke, 2017; Franke & Gaser, 2019). The current data set did not allow us to adapt a similar individualized approach and assess individual differences in aging trajectories. We encourage future work into this direction, particularly regarding our finding of increasing rich club organization throughout life. If increasing rich club organization was indeed a compensatory effort to maintain functional capacity as structural connectivity strength decreases, it should become particularly pronounced in individuals who age faster.

### Conclusion

We utilized the largest developmental sample with structural connectomes across the life span so far and applied non-linear statistical modeling to study life span trajectories of brain connectivity, network hubs, and rich club organization in the structural connectome. We confirmed ‘inverted U’ shaped trajectories for brain connectivity, found highly consistent network hubs across age groups, and found that rich club organization may remain relatively preserved in the aging brain. This might have implications for neural reserve and resilience in the aging brain and individual differences in biological and cognitive aging.

## Notes

### Competing Interest Statement

The authors have declared no competing interest.

