## Supplementary Material for "Trajectory of rich club properties in structural brain networks"

In the following, we are documenting additional analyses on network measures, different parameter choices,  subgroups, and alternative connection weights.

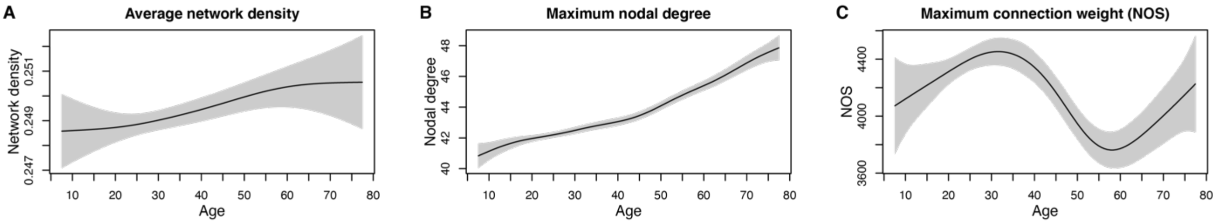


Supplementary figure 1: Average network density, maximum nodal degree and maximum connection weight (NOS) across the lifespan. Plotted are fitted values from GAMMs with 95% confidence intervals shaded in gray.

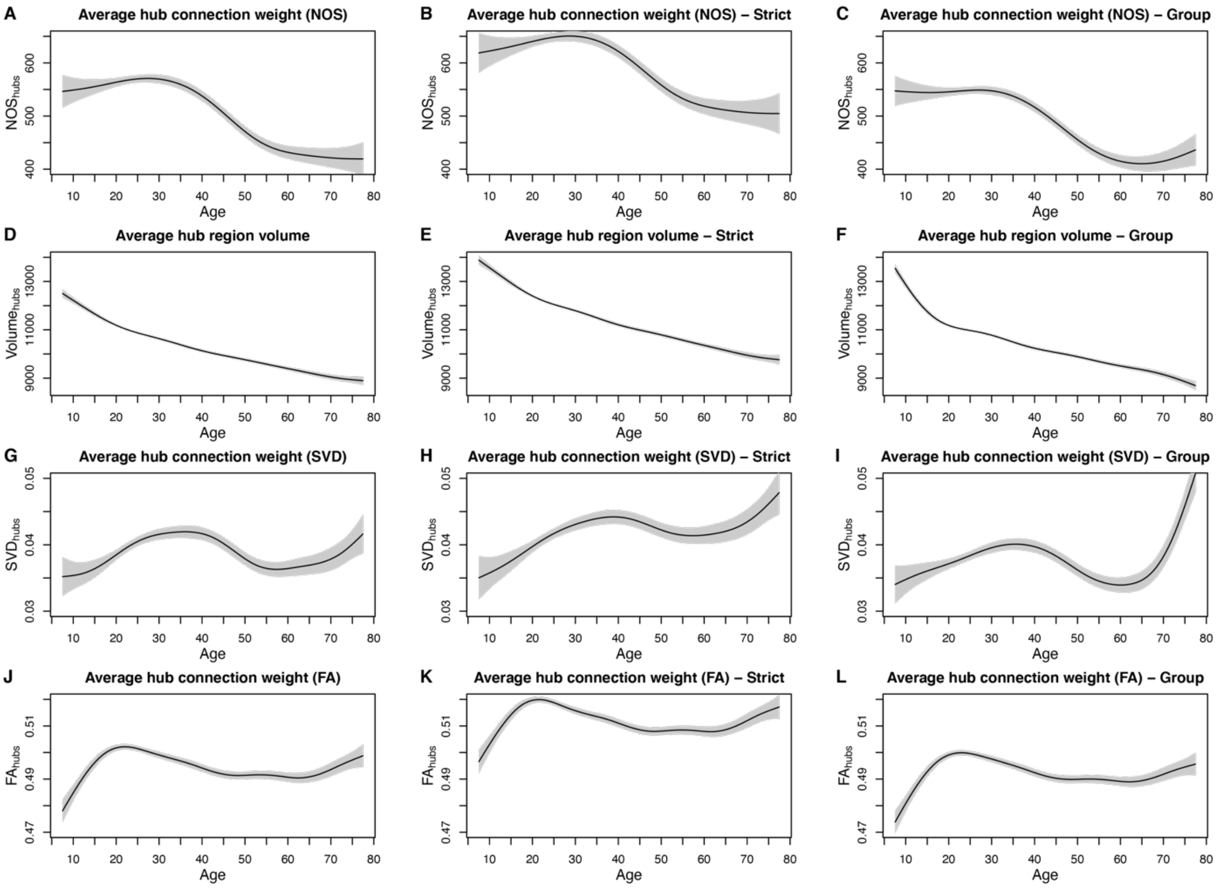


Supplementary figure 2: Average hub properties across the life span according to different hub partitions. Panels in rows correspond to: NOS-connection weights (row 1, A-C), regional gray matter volume (row 2, D-F), SVD-weighted connection weights (row 3, G-I) and FA-weighted connection weights (row 4, J-L). Panels in columns refer to hub partition according to the twelve brain regions that were most consistently identified as hubs across the entire sample (column 1), the nine brain regions that were identified as hubs in each age group (strict hub partition, column 2), and a group-specific hub definition of those twelve brain regions most consistently identified as hubs in the respective age groups (group hub partition, column 3). Plotted are fitted values from GAMMs with 95% confidence intervals shaded in gray.

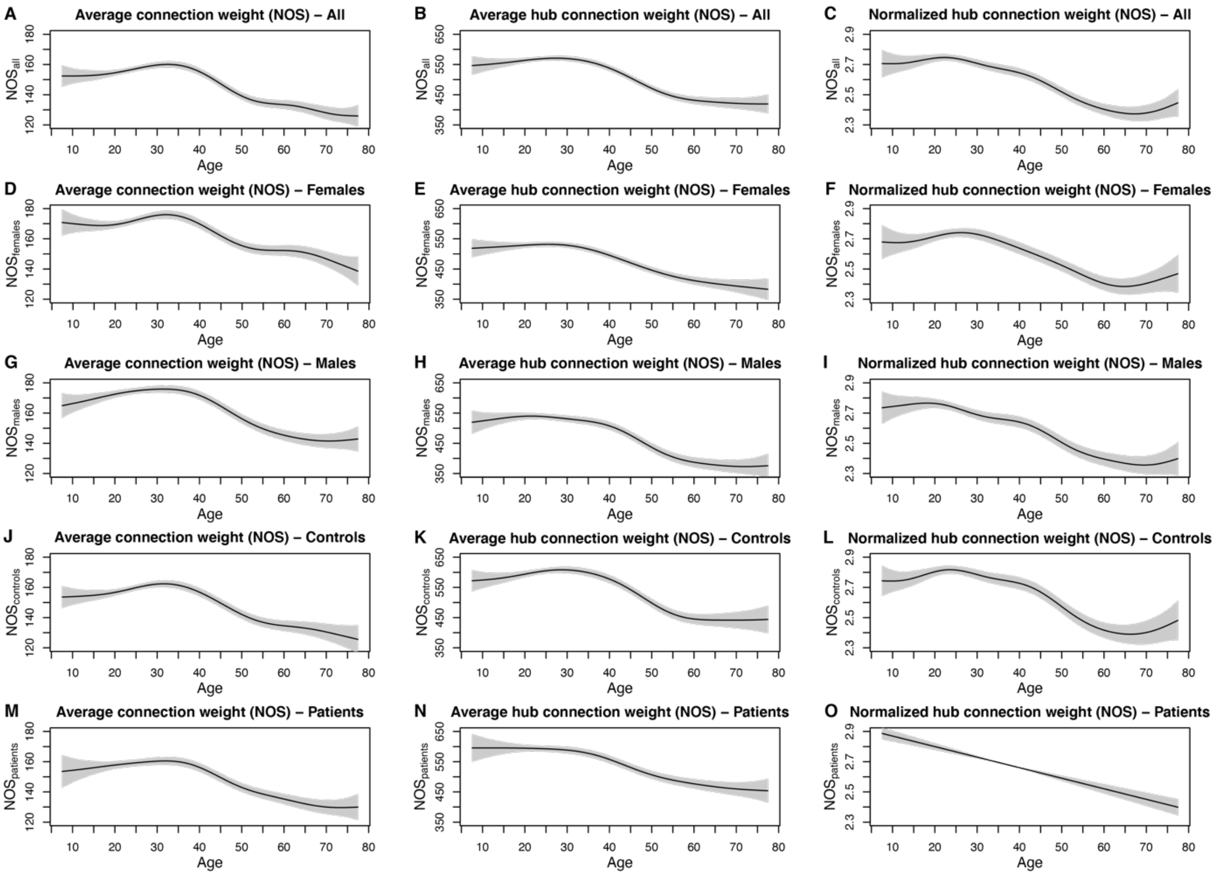


Supplementary figure 3: Average nodal and hub connection weights (NOS) across the life span including interactions of gender and disease status with age. Panels in rows correspond to different subpopulations: whole sample (row 1, A-C), females (row 2, D-F), males (row 3, G-I), controls (row 4, J-L), and patients (row 5, M-O). Panels in columns refer to averages across all brain regions (column 1), averages across all hub regions (column 2), and averages of hub regions relative to all brain regions (column 3). Plotted are fitted values from GAMMs with 95% confidence intervals shaded in gray.

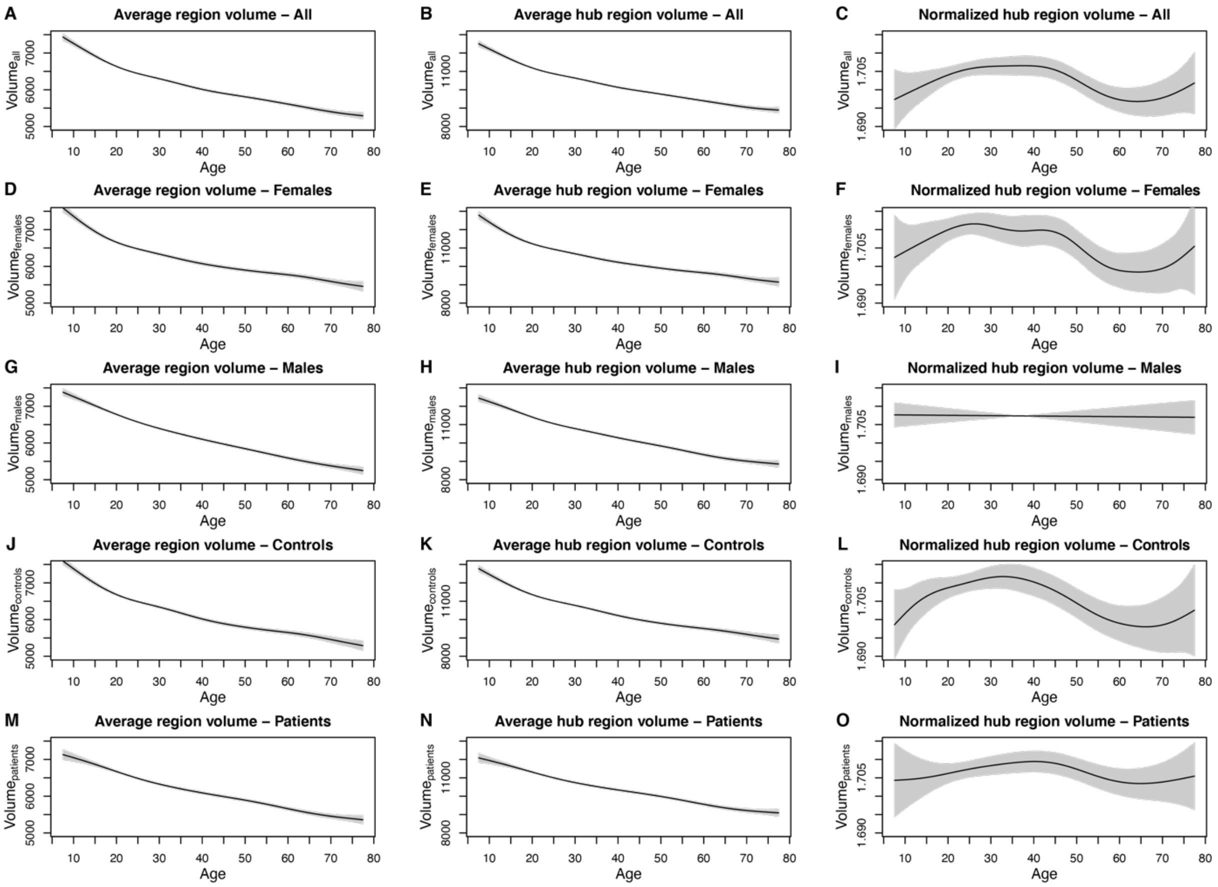


Supplementary figure 4: Average overall and hub regional gray matter volumes across the life span including interactions of gender and disease status with age. Panels in rows correspond to different subpopulations: whole sample (row 1, A-C), females (row 2, D-F), males (row 3, G-I), controls (row 4, J-L), and patients (row 5, M-O). Panels in columns refer to averages across all brain regions (column 1), averages across all hub regions (column 2), and averages of hub regions relative to all brain regions (column 3). Plotted are fitted values from GAMMs with 95% confidence intervals shaded in gray.

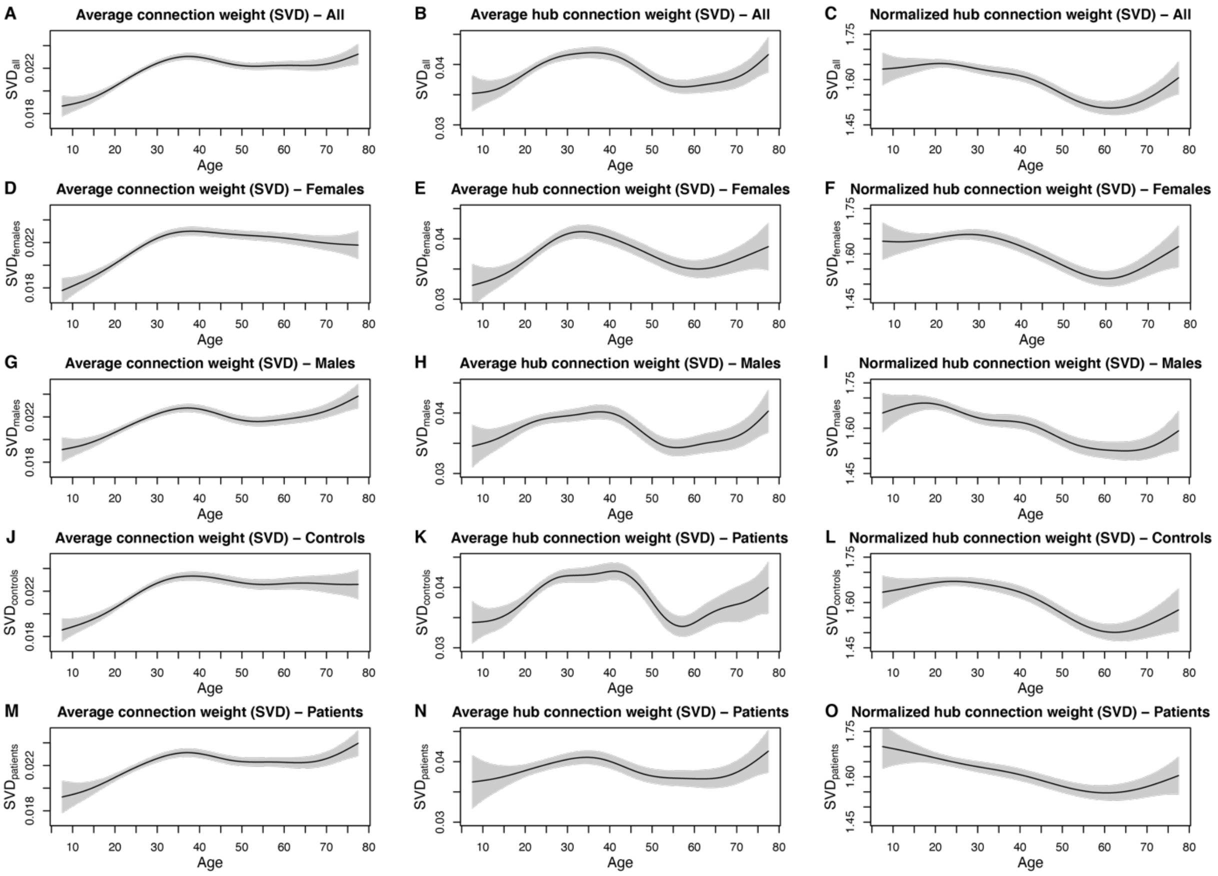


Supplementary figure 5: Average nodal and hub connection weights (SVD) across the life span including interactions of gender and disease status with age. Panels in rows correspond to different subpopulations: whole sample (row 1, A-C), females (row 2, D-F), males (row 3, G-I), controls (row 4, J-L), and patients (row 5, M-O). Panels in columns refer to averages across all brain regions (column 1), averages across all hub regions (column 2), and averages of hub regions relative to all brain regions (column 3). Plotted are fitted values from GAMMs with 95% confidence intervals shaded in gray.

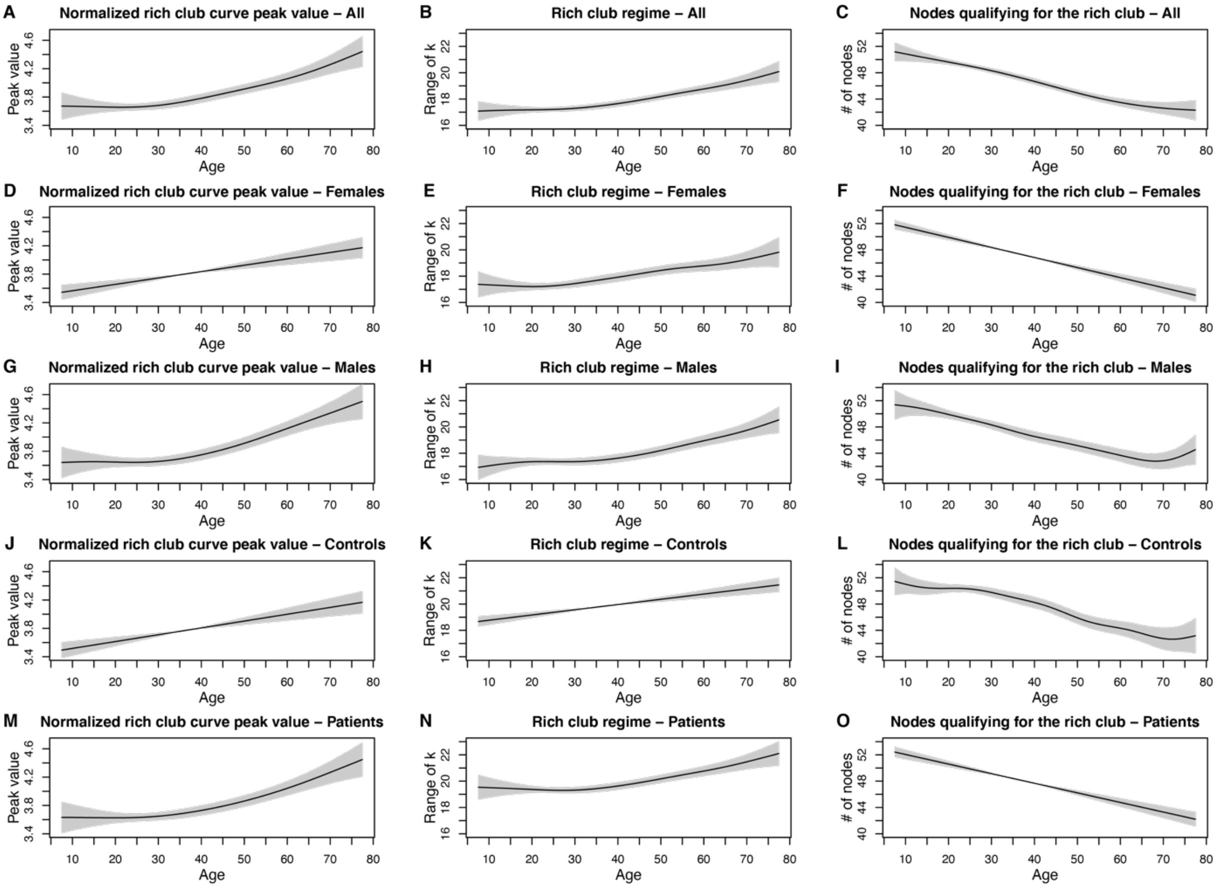


Supplementary figure 6: Average rich club property measures across the life span including interactions of gender and disease status with age. Panels in rows correspond to different subpopulations: whole sample (row 1, A-C), females (row 2, D-F), males (row 3, G-I), controls (row 4, J-L), and patients (row 5, M-O). Panels in columns refer to separate rich club measures: the maximum normalized rich club coefficient (column 1), the length of the range of consecutive degree levels k for which the empirical rich club curve exceeded the curve from the null models (column 2), and the number of nodes qualifying for rich club membership (column 3). Plotted are fitted values from GAMMs with 95% confidence intervals shaded in gray.

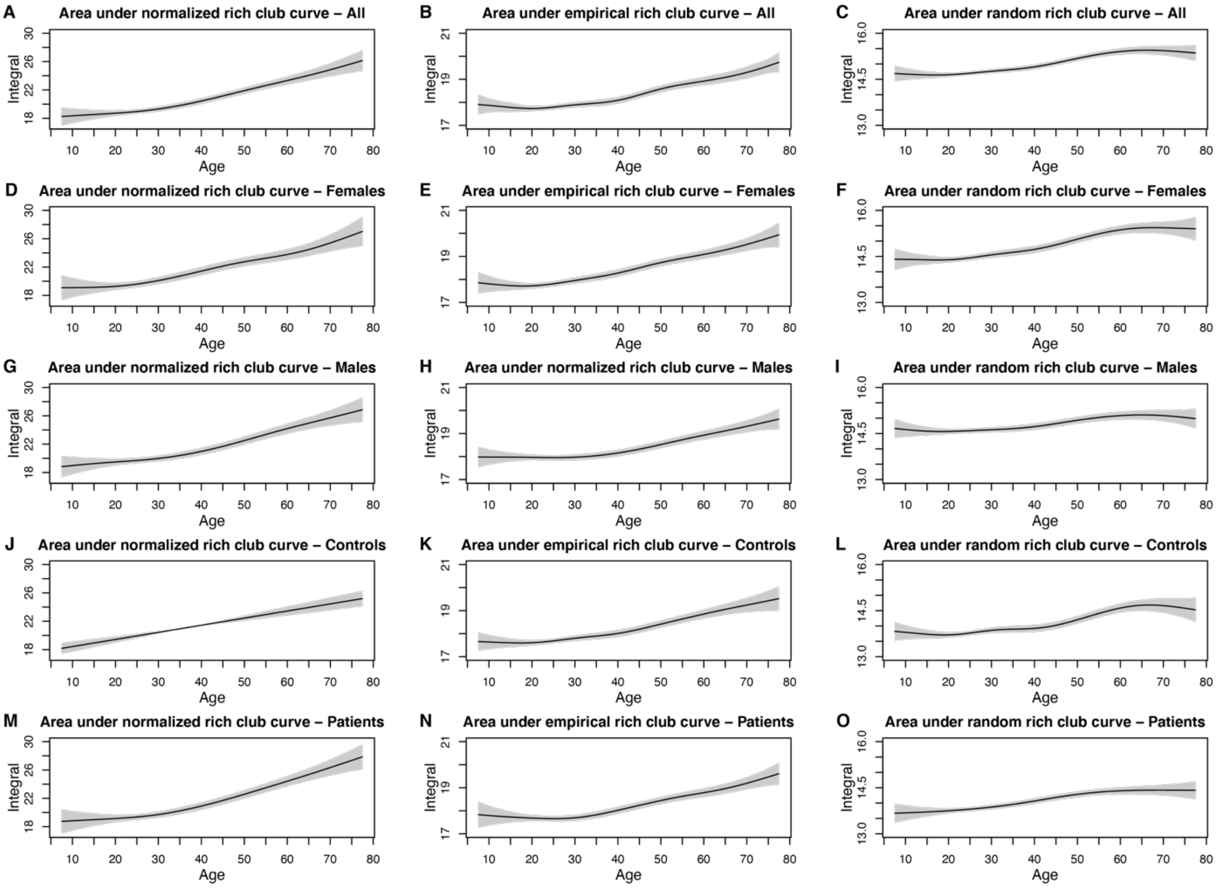


Supplementary figure 7: Average area under the rich club curves as a measure of the overall rich club property across the life span including interactions of gender and disease status with age. Panels in rows correspond to different subpopulations: whole sample (row 1, A-C), females (row 2, D-F), males (row 3, G-I), controls (row 4, J-L), and patients (row 5, M-O). Panels in columns refer to the area under the normalized rich club curve which refers to the overall strength of rich club organization in the network (column 1), and the areas under the rich club curve for the empirical networks (column 2) and the random null models (column 3). Plotted are fitted values from GAMMs with 95% confidence intervals shaded in gray.

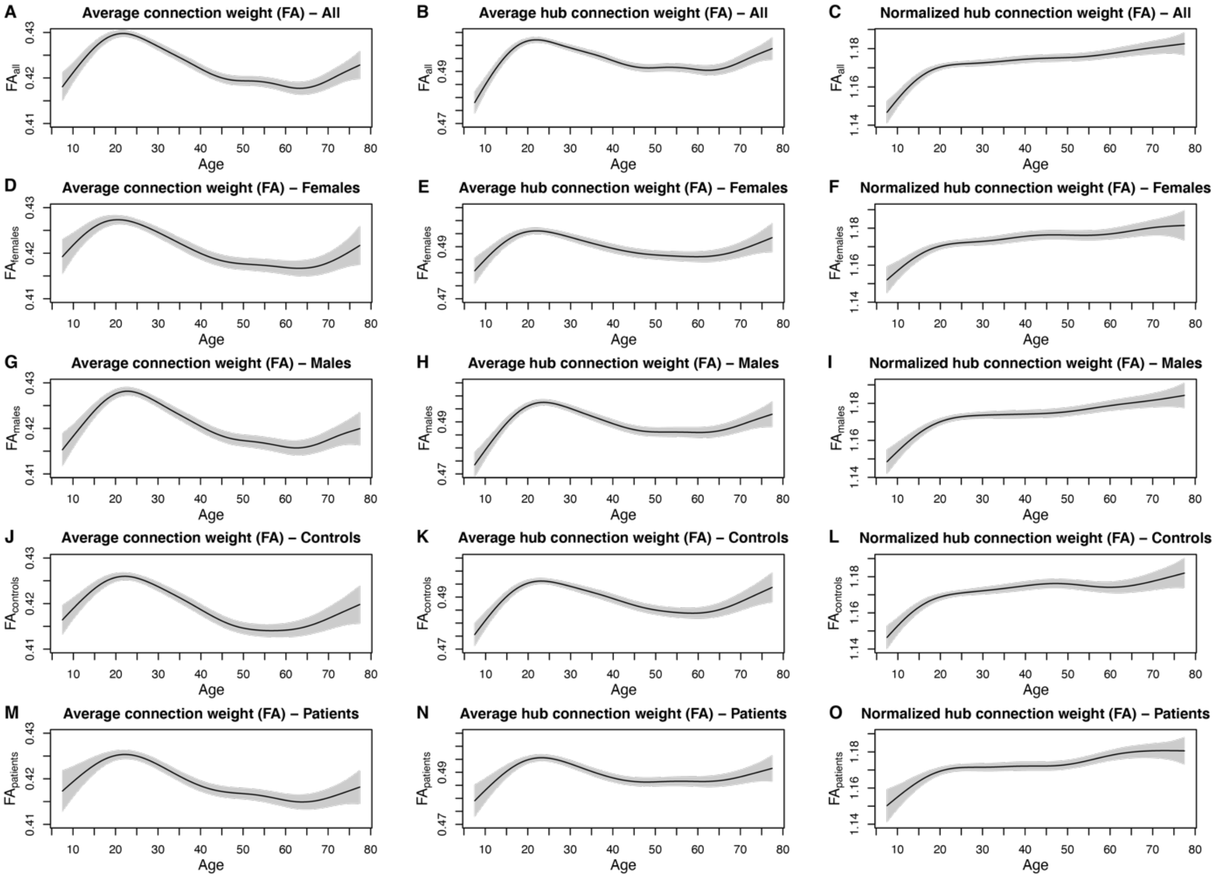


Supplementary figure 8: Average nodal and hub connection weight (FA) across the life span including interactions of gender and disease status with age. Panels in rows correspond to different subpopulations: whole sample (row 1, A-C), females (row 2, D-F), males (row 3, G-I), controls (row 4, J-L), and patients (row 5, M-O). Panels in columns refer to averages across all brain regions (column 1), averages across all hub regions (column 2), and averages of hub regions relative to all brain regions (column 3). Plotted are fitted values from GAMMs with 95% confidence intervals shaded in gray.
